# Phage-mediated lysis does not determine *Cutibacterium acnes* colonization on human skin

**DOI:** 10.1101/2025.09.09.675206

**Authors:** A. Delphine Tripp, Evan B. Qu, Ishaq Balogun, Julia Brodsky, Jacob S. Baker, Christopher P. Mancuso, Simon Roux, Fatima A. Hussain, Tami D. Lieberman

## Abstract

Despite *Cutibacterium acnes* being the most abundant and prevalent bacteria on human skin, only a single type of phage has been identified that infects this host. Here, we leverage this one-to-one system to systematically characterize how the phage-bacteria arms race shapes *C. acnes* evolution and community composition on individual people. Our analysis reveals a surprising lack of phage-mediated selection despite global prevalence of *C. acnes* phages. Analysis of anti-phage defense systems across 3,205 bacterial genomes revealed a limited, phylogenetically restricted defense repertoire under weak selective pressure to diversify or be maintained. Functional assays did not reveal alternative phage resistance mechanisms or fitness costs associated with defense gene carriage that could explain this limited immune arsenal. This lack of pressure to maintain phage resistance could not be explained by lack of phage colonization, as examination of 471 global human facial skin metagenomes demonstrated that even in samples with high virus-to-microbe ratio, phage-sensitive clades dominate on-person populations. Together, these findings indicate that phage pressure, while present, does not play a critical role in determining strain fitness and success within *C. acnes* populations on human skin. We propose that this observed weak phage-mediated selective pressure can be explained by the anatomy of skin: *C. acnes* growth is thought to occur at the bottom of pores, where exposure to phage may be limited by physical barriers. Together, this portrait of a static arms race provides a strong contrast with other microbial species in different ecosystems and expands understanding of phage-bacteria interactions in the human microbiome.

## Introduction

Understanding the selective forces that shape bacterial population structure in host-associated microbiomes is crucial for developing targeted microbiome-based therapeutics^1^,^2^. While extensive research has described strain-level diversity within human microbiomes^3^, the eco-evolutionary processes determining strain representation and success are unclear. There is growing theoretical and experimental evidence that phage-mediated selection is a likely driver of the abundance and distribution of resistant and sensitive bacterial strains^4–7^. Yet to date, the role of phage predation in shaping bacterial fitness, colonization success, and the overall population structure of commensal species remains poorly characterized.

Progress in understanding phage-mediated selection in host-associated ecosystems has been constrained by the lack of model systems that can bridge experimental validation of host ranges with strain-level tracking in natural populations. *Cutibacterium acnes (C. acnes)* on human sebaceous skin provides a tractable model to investigate how phage-mediated selection shapes bacterial population structure. *C. acnes* is a lipophilic, facultative anaerobe that primarily inhabits the sebum-rich pilosebaceous units of sebaceous skin and is the most abundant bacterial species on human skin^8,9^. On-person *C. acnes* populations arise from multiple colonization events, resulting in highly phylogenetically heterogenous, distinct strain types coexisting on individuals’ skin^10–15^ (Figure S1). This diversity is largely attributed to neutral ecological processes driven by the spatial architecture of the skin: physical segregation of bacterial subpopulations across pores reduces migration, competition, and ecological interactions, thereby promoting neutral coexistence^11^. Yet, much remains unknown about the potential role of phage predation in shaping *C. acnes* intraspecies diversity, and the impact of skin anatomy and physiology on the strength and outcomes of phage-bacteria interactions.

Intriguingly, despite the strain-level diversity found within *C. acnes*, only a single phage species has been isolated that can infect *C. acnes*^16–23^. This phage belongs to a single species of tailed phage (*Caudoviricetes* class) with a siphovirus morphology, which represents one of the most abundant viral groups in the skin microbiome^10,24–26^. *C. acnes* phages display striking genomic conservation, with pairwise ANI exceeding 85% across whole genomes of independently isolated phages^16–19,21–23^. These phage exhibits primarily lytic lifecycles, though they can be found occasionally in the cytoplasm of their host in a non-integrating, episomal prophage state known as a pseudolysogen^27^. Remarkably, host range assays have revealed broad and overlapping lytic activity, with all tested phages lysing 80-90% of tested clinical *C. acnes* isolates^16–19,21–23^, suggesting limited evolutionary diversification.

This simplified bacteria-phage system offers a unique advantage to study the impact of phage-mediated selection on bacterial population structure and examine if phage resistance confers fitness advantages that influence strain success on human skin. Understanding these dynamics has particular relevance given *C. acnes*’ role in both skin health and disease, the development of *C. acnes* as a probiotic chassis^28^, and the growing interest in phage-based therapeutic applications for acne^18,20,29–31^.

Here, we integrate whole-genome evolutionary reconstruction, experimental phage infectivity assays, and metagenomic analyses to study how encoding phage resistance influences *C. acnes* strain fitness and colonization success on human skin. We find that *C. acnes* maintains a limited arsenal of anti-phage defense systems, experiences weak selection to maintain these defenses, and shows pervasive signatures of inactivation in its defensome. Despite no detectable fitness costs associated with defense system carriage, we find that phage-sensitive clades represent the majority of *C*. acnes subpopulations in global facial skin metagenomes, even in communities with high viral loads. These findings suggest that phage resistance is not a major determinant of strain-level success *in vivo*, and confers minimal competitive advantages in the skin environment.

## Results

### The limited pan-immune arsenal of *C. acnes*

To understand the structure of phage-resistant and sensitive subpopulations on human skin, we began by profiling phage resistance in natural *C. acnes* isolates. We first characterized *C. acnes*’ pan-immune repertoire -- the complete set of anti-phage defense systems encoded across the species. We curated a comprehensive dataset of fully sequenced strains that represent all known *C. acnes* subspecies and major intraspecies clades (i.e., phylogroups and subphylogroups). Our collection comprised 3,205 genomes, including 173 isolates annotated as ‘*Cutibacterium acnes*’ in the 661K bacterial genome database^38^ and 3,032 isolates previously sampled by our lab from the sebaceous skin of 43 subjects^11,12^ (Figure S1; Table S1).

Scanning for known anti-phage defense systems using DefenseFinder^32^ revealed that *C. acnes* maintains a very limited immune arsenal. Individual genomes carry a maximum of four defense systems, with more than 40% of genomes lacking detectable defenses altogether (range 0 to 4; Table S2). This limited arsenal is typical of the genus *Cutibacterium*, which has been documented to encode the smallest pan-immune repertoire within the phylum *Actinobacteria*^53^. This limited arsenal is significantly lower than what has been detected in genomes from the genera *Corynebacterium, Staphylococcus* and *Streptococcus*, which are associated with human skin and have similar sized genomes, but larger pangenomes^54^ (Figure S2).

The entire *C. acnes* pan-immune system consists of only six distinct elements: a novelly identified Gabija system and previously described CRISPR Cas Type I-E, Lamassu, Restriction-Modification (RM) Type III, RM Type IIG, and an Abortive Infection (AbiD) system^55–57^. These systems are co-localized across three genomic islands present in phylogroups D, H, and K (Figure 1A). Defense islands are carried by 100% of isolates within these phylogroups (Figure S3; Table S2), resulting in considerable homogeneity in defenses among genomes at both coarse- and fine-grained phylogenetic resolutions.

**Figure 1.**
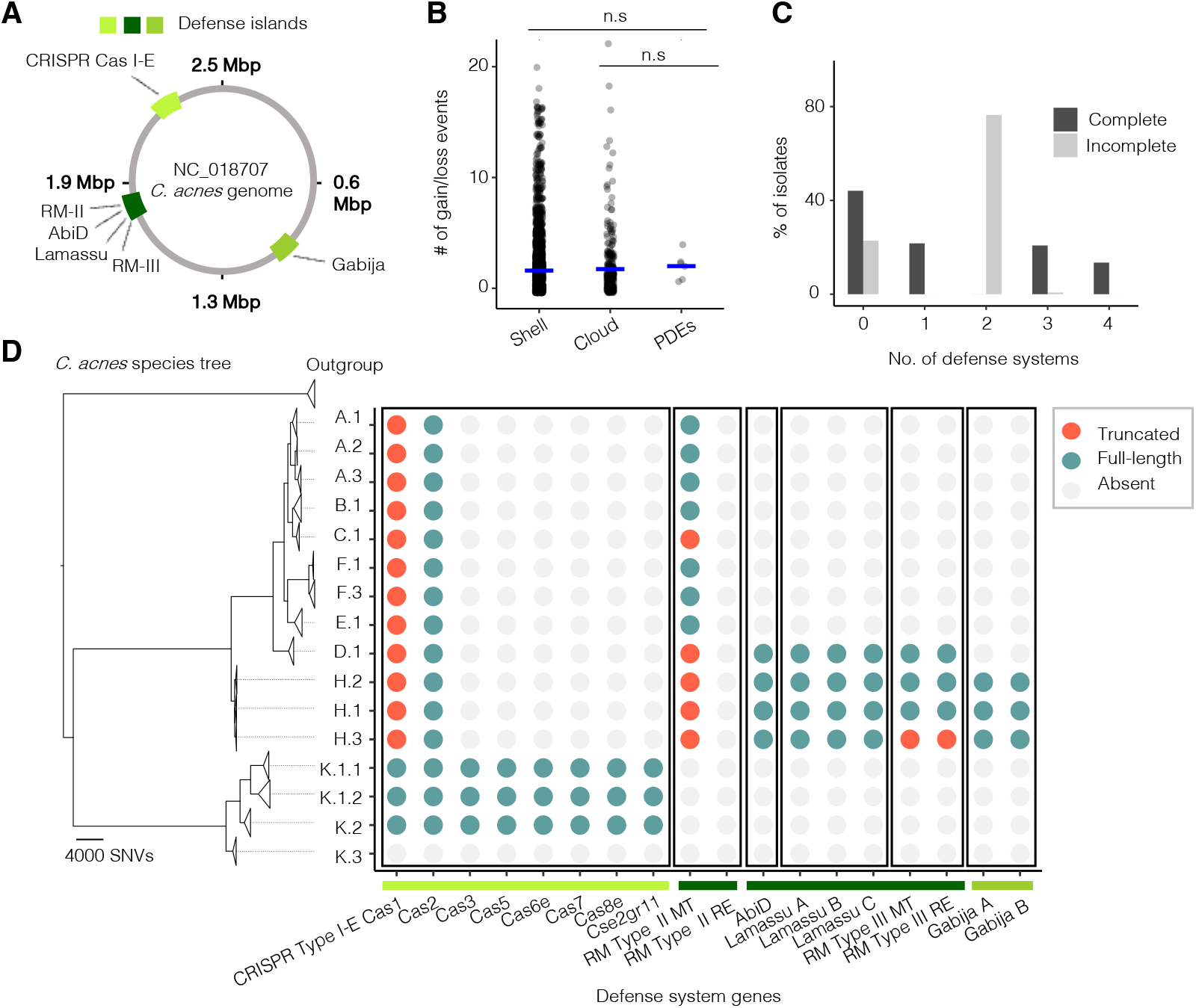
Weak selection to maintain or diversify the limited immune arsenal of *C. acnes*. (A) The *C*. acnes pan-immune system consists of six distinct defense systems that co-localize onto three defense islands present in phylogroups D, H, and K (Table S2; Methods). (B) Ancestral trait reconstruction of the defensome and accessory genome (shell and cloud) inferred using pastML (Methods) shows no significant difference in the number of estimated gain/loss events. (C) Bar chart of the number of complete or incomplete defense systems (e.g. with missing or truncated mandatory genes; Methods) reported per genome versus the percentage of genomes. (D) A maximum-likelihood phylogenetic tree constructed at the subphylogroup level and truncation analysis reveal pervasive signatures of defense gene loss through historical inactivation. Individual genes are colored in each genome by whether or not the gene is present in full, absent, or truncated, and genes are grouped by defense islands (bottom bars).

### Lack of diversification and rapid turnover in *C. acnes* defense genes

We tested if the limited arsenal of defense systems in *C. acnes* could be explained by genome-wide evolutionary constraints limiting acquisition of new defenses. *C. acnes* is known to exhibit low rates of horizontal gene transfer^14,55,58,59^. If this low species-level rate of gene flux was solely responsible for *C. acnes’* limited arsenal, we would expect defense genes to be exchanged at disproportionately higher rates than other accessory genes^60^, given strong selective pressure for phage resistance. To quantify gene turnover, we determined homologous gene groups and used ancestral trait reconstruction to estimate the total number of gain and loss events for each gene over *C. acnes* evolutionary history (Methods). We found no significant difference in the number of gain and loss events between defense and accessory genes (p = 0.09 for the shell, and p = 0.09 for the cloud; Figure 1B), indicating that the defensome evolves at the same rate as the rest of the accessory genome.

Intriguingly, many non-defense genes exhibited higher rates of gain and loss compared to those in the defensome. This includes genes from *C. acnes* prophages -- maintained as non-integrated, mobile pseudolysogens -- and two distinct plasmids that, despite being chromosomally integrated, display extensive HGT across isolates: a linear tight adherence (TadE) plasmid^61–64^ and a novelly identified circular rep-encoding single-stranded (pCRESS) plasmid^65^ (Figure S3). These results highlight that the conserved nature of the *C. acnes* defensome is not due solely to genome-wide constraints on gene flux.

### Weak selection to maintain the immune repertoire of *C. acnes*

The absence of rapid evolution in defenses may indicate a lack of strong selection to maintain protein-encoded phage resistance in *C. acnes*. We reasoned that weak selection on anti-phage defense systems might result in systematic defense degradation through pseudogenization and deletion events in the evolutionary history of the *C. acnes* pan-immune system. To test this hypothesis, we screened for full-length and truncated variants of defense genes (Methods) and analyzed their presence and absence in the context of the core genome phylogeny. Notably, 12 out of 16 subphylogroups, nearly 80% of isolates, carried one or more incomplete defense systems (Figure 1C). Here, incomplete refers to defense systems with one or more core genes deleted or disrupted by frameshifts or premature stop codons.

Evolutionary inference suggests that these incomplete defense systems likely arose from inactivation events that occurred early in the speciation of *C. acnes* (Figure 1D). We found evidence of both historical (i.e., CRISPR Cas Type I-E in phylogroups A - H; RM Type II in phylogroups C and D - H) and recent (RM Type III in subphylogroup H.3) defense gene loss and truncations. Notably, some defense systems were likely lost multiple times, as previously described for the CRISPR Cas Type I-E and the RM Type II systems^56–58^. These pervasive signatures of inactivation across multiple defense systems, in combination with a conspicuous absence of evidence for recent system gains (most recent gain was the Gabija system in the MRCA of phylogroup H), provides compelling evidence of weak selection to maintain protein-encoded phage resistance in *C. acnes*.

Collectively, these findings suggest that the widespread lack of robust defenses against phage within *C. acnes* likely arise from weak selection to acquire and maintain defenses, rather than from overall genome content inflexibility.

### Defense systems are biologically active and the primary determinant of phage resistance

Weak selection to maintain protein-encoded phage resistance could be explained by alternative resistance mechanisms being more common in natural isolates than previously recognized. Beyond protein-encoded defenses, bacteria can intrinsically acquire resistance to phage infection through multiple strategies^66,67^, including superinfection exclusion, and *de novo* mutations affecting phage attachment or entry. We ruled out superinfection exclusion as a common strategy for phage resistance in nature, as only 3% of 3,205 sequenced *C. acnes* isolates carried prophages (Figure S3). We note that the true rate of prophage carriage may be even lower than estimated here by the prevalence of pseudolysogens, because more than half of pseudolysogen-containing isolates (57.1%) originated from just two subjects, suggesting high inter-individual variation or potential sample bias (Figure S4).

To functionally screen for phage resistance, we curated a library of 109 isolates representing *C. acnes* diversity at multiple evolutionary scales (Figure S5), and then experimentally tested bacterial susceptibility to phage. This library included sets of closely related isolates from the same lineage (separated by fewer than 100 SNVs across their whole genomes) to capture fine-scale evolutionary changes in phage resistance profiles. We assessed susceptibility to phage by growing each isolate with and without a phage cocktail in liquid culture and calculating growth inhibition scores based on OD_600_ (Methods; Table S3).

We first evaluated the relative importance of defense systems versus alternative resistance mechanisms in conferring resistance. Isolates carrying defense systems were inhibited significantly less compared to those lacking defenses (mean inhibition score of 18% vs. 68.6%, respectively; p < 0.001; Figure 2A and S6). Notably, the highest observed inhibition score was 84.2%, as some bacterial growth always occurred before phage lysis initiated. Most phylogroups exhibited low variability in inhibition scores (mean absolute deviation, MAD range 0.024 to 0.071; Figure 2A), with the exception of phylogroup K (MAD = 0.234). This phylogroup encodes a CRISPR Cas Type I-E system known to confer spacer-dependent anti-phage activity^16,17^. Indeed, analysis of phylogroup K CRISPR Cas array spacers confirmed that the high within-phylogroup variability in inhibition scores reflects the phylogenetic distribution of CRISPR Cas spacers: spacer content is highly conserved within lineages but diverges at broader phylogenetic scales (Figure S7; Table S4).

**Figure 2.**
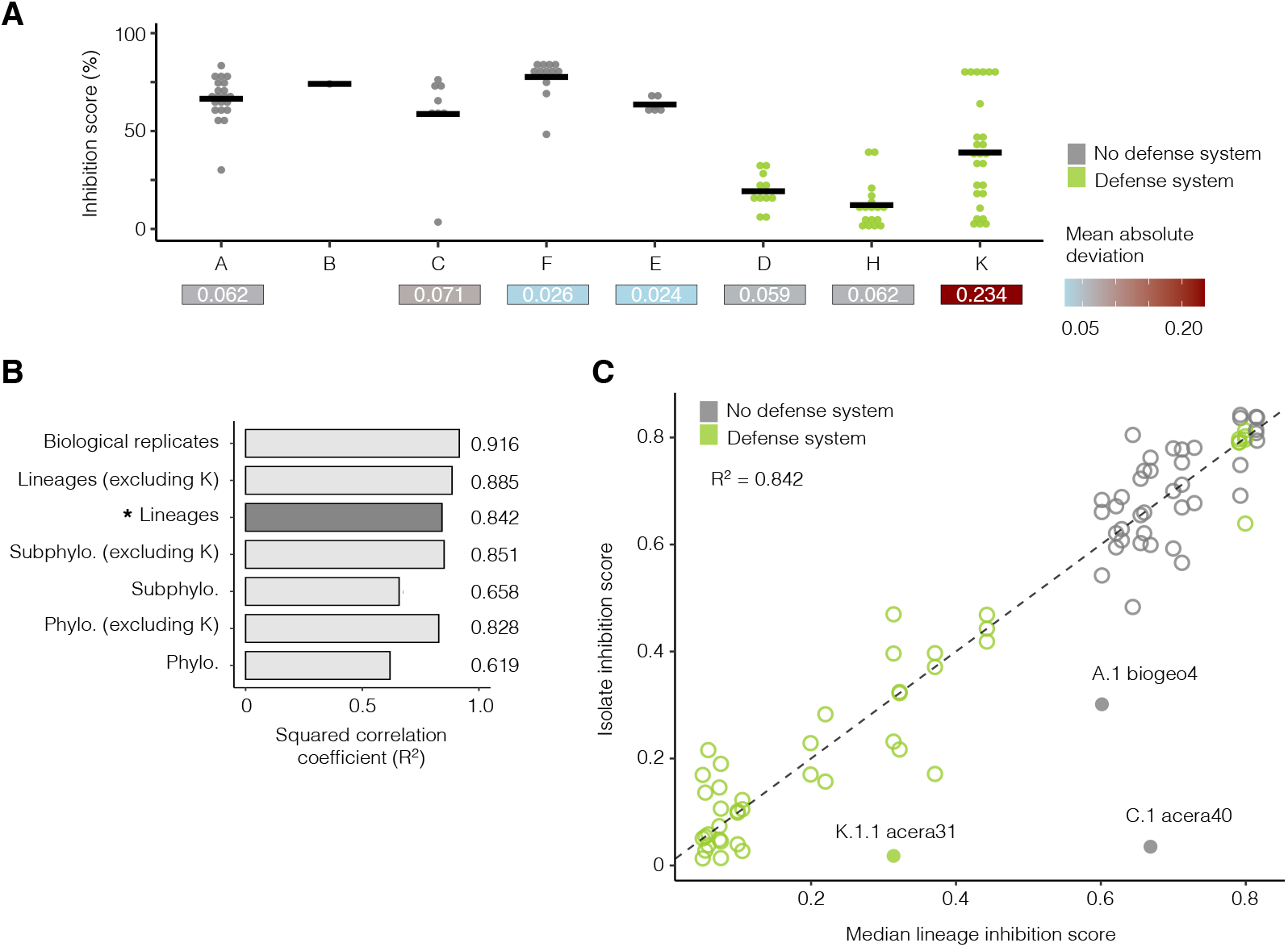
Phage resistance profiles are highly conserved and predictable from coarse phylogenetic groupings. (A) Isolates lacking defense systems (grey) show significantly higher inhibition scores compared to those carrying defense systems (green), indicating increased susceptibility to phage (Figure S6). Inhibition scores are calculated as the ratio of the area under the curve (AUC) for untreated versus phage-treated bacterial cultures (Methods). Notably, the highest observed inhibition score was 84.2%, as some bacterial growth always occurred before phage lysis initiated. Within-phylogroup variation is particularly high among isolates from phylogroup K, which encode a CRISPR Cas Type I-E system known to confer spacer-dependent anti-phage activity (Figure S7). (B) Bar chart showing the squared correlation coefficient (R^2^) between each isolate’s inhibition score and the median score of its phylogenetic group at the lineage, subphylogroup, and phylogroup levels, with and without phylogroup K. High R^2^ values across all taxonomic levels indicate that resistance profiles are strongly conserved and largely static, with minimal variation even at broader evolutionary scales (Figure S8 and S9). (C) Scatter plot of individual inhibition scores versus the median inhibition score for all isolates within the same lineage. The dashed line represents y = x. Points are colored by defense system status (grey: absent; green: present), with outliers highlighted as closed circles.

### Phage resistance profiles are highly conserved and predictable from coarse phylogenetic groupings

To screen for phenotypic variation in bacterial susceptibility to phage beyond the phylogroup level, where defenses are heavily conserved, we analyzed inhibition score divergence across phylogenetic scales (lineage, subphylogroup, phylogroup) using pairwise distance comparisons. No significant differences in pairwise distances were detected among taxonomic levels (all comparisons p > 0.1; Figure S8), indicating that the rate of phenotypic divergence in inhibition scores remains consistent across evolutionary scales. To evaluate whether this uniformity reflects evolutionary stasis or masks rapid phenotypic turnover, we tested the phylogenetic conservation of inhibition scores by correlating each isolate’s score with the median of its respective phylogenetic group at the lineage, subphylogroup, and phylogroup levels. High correlations were observed at all levels (Figure 2B), approaching the reproducibility of biological replicates, with conservation strengthening at finer taxonomic resolutions (Figure S9). Notably, inhibition score conservation at the lineage-level exceeded that of bacterial growth rates (r^2^ = 0.84 versus 0.69; Figure S9), suggesting that in *C. acnes* susceptibility to phage is more tightly linked to evolutionary history than general physiological traits.

We observed two exceptions to this trend -- isolates that lacked defense systems yet exhibited low inhibition scores -- providing rare evidence of alternative resistance mechanisms (Figure 2C). In one case, resistance evolved *in vitro* at hour 24 of the experiment (Isolate A.1 biogeo4; Figure S10). The other represents the sole observed case of naturally occurring resistance evolved *in vivo* (Isolate C.1 acera40; Figure S10). In both cases, two other isolates from these lineages, sampled from the same subject at the same timepoint, remained susceptible to phage infection. Overall, these exceptions highlight that while *de novo* resistance can emerge in lab culture or on human skin, these spontaneous resistance strategies are uncommon in natural populations. Consequently, defense systems represent the major determinant of anti-phage activity in *C. acnes*, and resistance profiles can be accurately predicted from coarse phylogenetic groupings.

### Phage-sensitive strains are enriched in global facial skin metagenomes

To assess the impact of defense system carriage on bacterial fitness *in vivo*, we turned to publicly available metagenomic data. We leveraged the fact that resistance profiles can be predicted from coarse phylogenetic groupings to estimate the proportion of each person’s *C. acnes* population that is composed of resistant and sensitive strains. To capture global populations, we analyzed 471 publicly available human facial skin metagenomes from individuals across the USA, Europe, and China, with each geographical region represented by samples from at least two independent research groups (Figure 3A and S11-13; Table S5).

**Figure 3.**
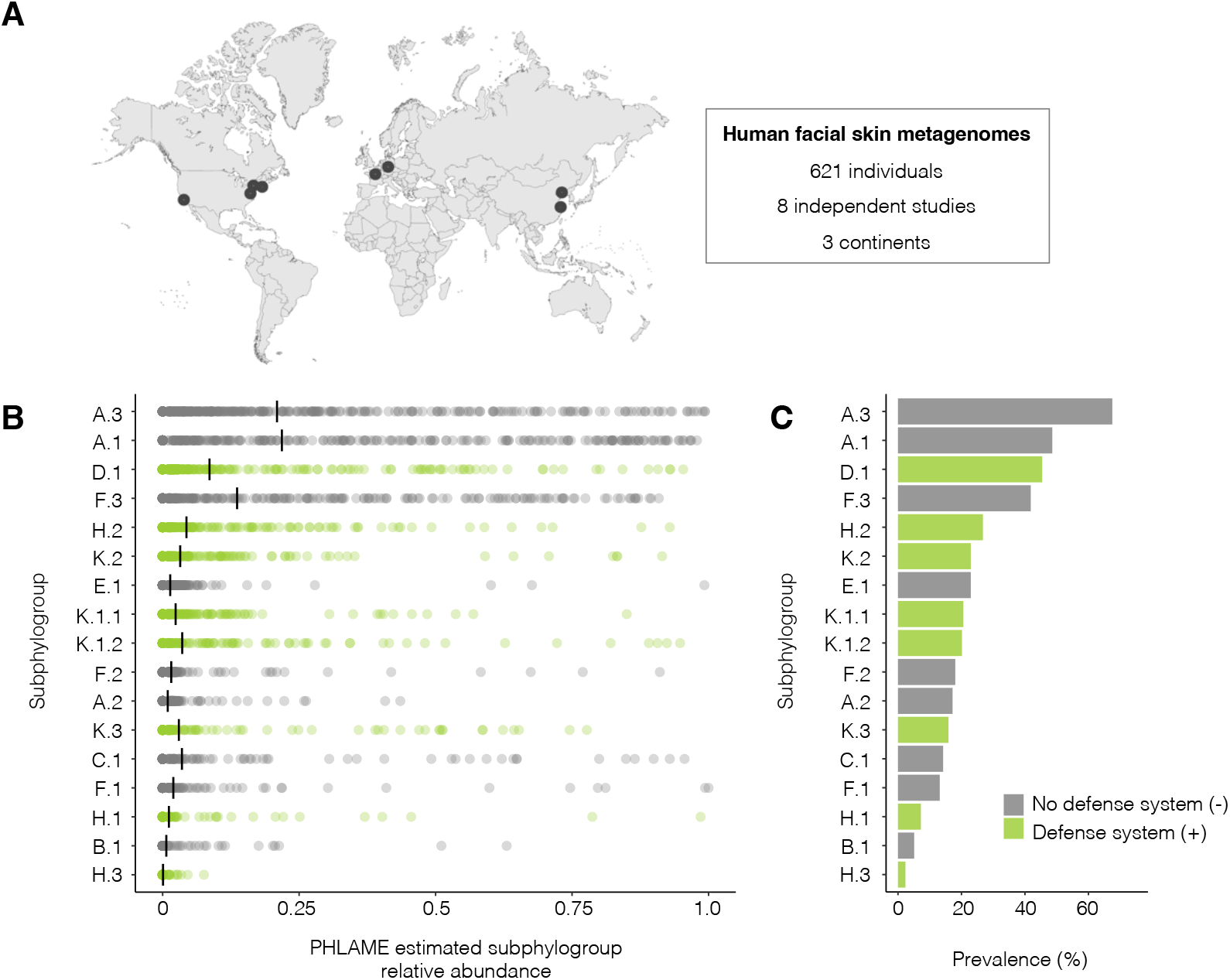
Phage-sensitive strains are enriched in global facial skin metagenomes. (A) We analyzed 621 publicly available human facial skin metagenomes from 3 continents across 8 independent studies originating in the USA, Europe and China. Each region contained data from at least two independent research groups. After quality control filtering (Methods), the final dataset consisted of metagenomes from 471 distinct individuals (Table S5; Figure S11). We used PHLAME (Methods) to estimate *C. acnes* subphylogroup (B) relative abundance and (C) prevalence in the 89.6% (422/471; Figure S12) of samples which had at least 5X average reference genome coverage (NC_018787). The most prevalent subphylogroups were A.3 and A.1, which lack complete defense systems. Generally, subphylogroups that lack defenses were found at significantly higher relative abundances (p < 0.001) amongst on-person *C. acnes* populations.

We determined the presence and relative abundance of each *C. acnes* subphylogroup using PHLAME^42^ (Methods), which uses core genome mutations to classify intraspecies diversity, and then inferred the total proportion of phage-resistant to sensitive strains. Across global facial metagenomes, phage-sensitive subphylogroups were detected at significantly higher relative abundances compared to resistant subphylogroups (p < 0.001; Figure 3B), with 3 of the top 4 most prevalent subphylogroups lacking defenses (Figure 3C). Together, phage-sensitive phylogroups dominate *C. acnes* populations on human facial skin, making up on average 70.4% of a person’s *C. acnes* population -- a ratio of 2.3 sensitive cells to every 1 resistant cell.

To test if the success of the sensitive strains in the population could be due to fitness costs imposed by carrying anti-phage defense genes, we assessed *C. acnes* growth rates *in vitro*. We observed no significant differences between defense-carrying and non-defense isolates (p > 0.1; Figure S6). While we cannot rule out tradeoffs that are difficult to detect *in vitro*, these results suggest that fitness tradeoffs alone are unlikely to explain the predominance of sensitive *C. acnes* subpopulations.

### Low *C. acnes* virus-to-microbe ratios in global facial skin metagenomes

We next investigated if the predominance of sensitive *C. acnes* subpopulations could be explained by low phage prevalence on human skin. Prior metagenomic surveys have hinted that *C. acnes* phage are common members of the human skin virome^26^. In our dataset, we indeed find that both phage and bacterial are common across geographies, with 59.7% (281/471) of individuals carrying *C. acnes* phage and 89.6% (422/471) carrying the bacteria (Methods, Figure S12). Among individuals carrying *C. acnes* bacteria, 65.2% (275/422) also harbored *C. acnes* phages, indicating frequent co-occurrence of host and phage populations.

To see if sensitive strains were more likely to persist on individuals without significant phage populations, we approximated the relative abundances of *C. acnes* virus and microbial densities using metagenome-derived virus-to-microbe ratios (mVMRs). When calculating VMR from metagenomes, it is crucial to consider that high frequencies of lysogens (dormant viruses integrated into bacterial genomes) can confound the assumption that viral genomes are solely derived from free virions^68–71^. *C. acnes* provides a distinct advantage, due to the absence of lysogens and low prevalence of pseudolysogens (detected in <3% of isolates, Figure S4) in this species. Thus, confounding by lysogens is likely negligible and the calculated mVMR more accurately reflects the true ratio of free viral to bacterial genomes.

Across individuals, mVMRs were strikingly low (median 0.04; range 0–15.6; Figure S13), corresponding to ∼1 viral genome per 25 bacterial genomes. This ratio contrasts with aquatic ecosystems, where viral genomes typically outnumber bacterial genomes by 10:1 or more^69^. In the human gut, metagenomic estimates suggest an mVMR of ∼4:1, though this ratio is likely inflated by sequencing of dormant lysogens, which dominate viral reads^68^. Direct quantification of virus-like particles and bacterial cells in gut samples yields a VMR of ∼0.1^68^, which aligns closely with our observations on skin. In the gut, this low rate of active lysis is thought to emerge from superinfection protection provided by carried lysogens. Since *C. acnes’* suppression of lysis via pseudolysogen-mediated superinfection exclusion is likely rare, its low mVMR suggests that phage lytic activity may be intrinsically limited on human skin.

### No competitive advantage for phage-resistant strains even in high viral load environments

We examined the relationship between VMR and the proportion of phage-sensitive to resistant strains across individuals. If phage exert strong selective pressure, we would expect resistant strains to be enriched in metagenomes with higher viral loads due to conferred fitness advantages. Instead, we observed no significant correlation between the estimated total relative abundance of resistant subphylogroups and the mVMR (p = 0.41; Figure 4A), with high mVMR samples spanning the full range of resistant subphylogroup relative abundances. These results illustrate that even in environments with high viral loads, phage resistance does not confer a strong competitive advantage on human skin at the community level.

**Figure 4.**
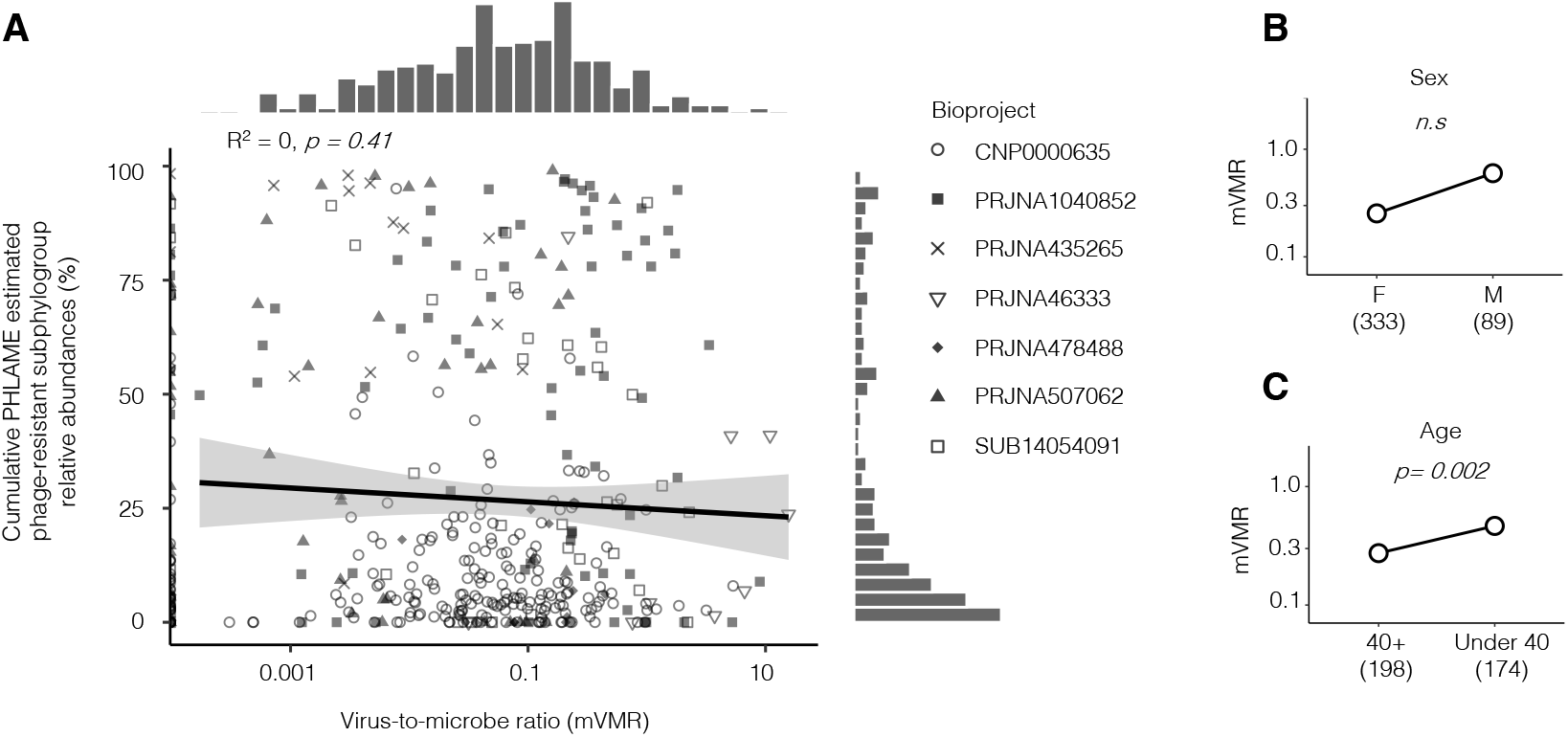
No competitive advantage for phage-resistant strains even in high viral load environments. (A) Scatter plot of the summed frequency of phage-resistant subphylogroups (strains carrying anti-phage defense systems) versus the mVMR (Methods). There is no significant correlation (p > 0.1). A histogram of each variable alone is shown on the top and the side. Shapes represent different studies. In the majority of samples, the mVMR was low – approximately 1 phage genome per 25 bacterial genomes (Figure S13). (B-C) Comparison of mVMR with human host demographic data reveals no association with biological sex (p > 0.1), but a significant association with subject age (p < 0.001). Age is stratified to a binary classification because continuous ages are not available for all subjects (Table S5).

We noted that a small portion (∼6%) of metagenomes exhibited mVMRs greater than or equal to one, prompting us to investigate whether this variation is related to human host factors. To limit confounding factors, we examined demographics that were consistently reported across multiple studies: biological sex and age. While biological sex did not significantly influence mVMR (p = 0.01), individuals under 40 years of age displayed significantly higher mVMRs compared to older individuals (p = 0.002; Figure 4B-C). This finding suggests that age-related shifts in skin physiology or strain composition may modulate virus-microbe dynamics independent of resistance traits.

Together, our results reveal that phage-mediated lysis does not play a major role in determining the success of *C. acnes* strains on human skin. The widespread presence of *C. acnes* phages across global populations, coupled with the predominance of phage-sensitive strains even in high-viral environments, suggests that other factors -- including skin spatial architecture, host immune interactions, nutrient competition, priority effects, antimicrobial resistance, or interbacterial warfare -- may be more important drivers of *C. acnes* evolution and population structure.

## Discussion

Bacteria-phage coevolutionary dynamics are typically characterized by an arms race, wherein defense system acquisition and counter-adaptations drive rapid antagonistic coevolution^60,72,73^. In this work, we demonstrate that the dominant skin commensal, *C. acnes*, is in an evolutionary static equilibrium with its only known phage. Our findings reveal a population dominated by phage-sensitive bacteria (Figure 3), characterized by a limited immune repertoire with widespread defense system inactivation (Figure 1), and no detectable fitness advantage for resistant strains, even in phage-rich environments (Figures 4). Despite retaining biologically active defense systems (Figure 2), their evolutionary stasis and phylogenetic conservation starkly contrast the rapid turnover expected under canonical Red Queen dynamics^5^. This static equilibrium suggests that ecological constraints within the human skin ecosystem can override the evolutionary forces driving classical virus–microbe arms races.

A key question arises: why is selective pressure not sufficient to drive the evolution of *de novo* resistance mechanisms in *C. acnes* populations, even when phage are present? Along these lines, why is there no strong selective pressure to maintain anti-phage defense systems despite experimental validation of their ability to confer resistance to phage? We posit two non-exclusive explanations: (1) physical barriers limit phage-bacteria encounters *in vivo*, and (2) skin conditions may induce phage-resistant physiological states.

We hypothesize that the spatial segregation of *C. acnes* within pilosebaceous units likely imposes a fundamental constraint on phage-mediated selection. Analogous to island biogeography, pores function as isolated microbial habitats where limited bacterial migration and phage dispersal may reduce encounter rates. Theoretical models predict that such fragmentation diminishes phage effective population sizes, restricting opportunities for coevolutionary escalation^74^. Even when lysis occurs within a pore, virions may struggle to disperse and infect neighboring follicles, creating localized “hotspots” of phage activity that cannot drive population-wide selection. This aligns with our observations of low metagenomic virus-to-microbe ratios (Figure S13) and evolutionarily static phage resistance profiles (Figure S9). Furthermore, the rarity of pseudolysogens (Figure S4) provides supporting evidence of limited phage dispersal, as superinfection exclusion -- a key driver of lysogeny in other systems -- is virtually absent. Future spatially resolved metagenomics (e.g., pore-level microsampling) will be needed to test whether phage activity varies microbiogeographically, reflecting these predicted hotspots and patch-level population dynamics amongst individual pores.

Beyond spatial barriers, skin physiology may modulate phage infectivity and efficacy. *C. acnes* is thought to replicate primarily at the base of follicles, where anaerobic, lipid-rich conditions differ starkly from the aerobic skin surface. While our *in vitro* assays confirmed phage activity under standard growth conditions, transcriptional dormancy in low pH, sebum-rich environments could reduce bacterial susceptibility to phage on human skin. A recent metatranscriptomic study reported that only a low proportion of *C. acnes* cells are transcriptionally active on skin^75^. This low transcriptional activity suggests that most *C. acnes* subpopulations are likely dormant and thus in a phage-resistance physiological state. For this reason, we encourage future investigation of phage-bacteria interactions under growth conditions that mimic the physiology of human skin to better understand *in vivo* dynamics.

Notably, the unknown forces that weaken phage-mediated selective pressure on *C. acnes* remain insufficient to completely eliminate its phages from skin populations. The coexistence of phage, resistant, and sensitive strains within individuals suggests that there are microenvironments on human skin where phage infection is successful. Furthermore, the lower abundances of phage on older individuals suggests that the relative abundance of this microenvironment can shift over time. Future work will be needed to determine whether these microenvironments are spatial or temporal.

One limitation of our study is that our conclusions hinge on the assumption that cultivated *C. acnes* phage represent the full diversity of natural populations. Although metagenomic and culturomic studies have not identified any novel *C. acnes* phage clades to date^10,16,17,19,24–26^, uncultivated viruses could potentially employ alternative infection strategies that evade detection. For example, the non-lytic chronic infection life cycle of certain ssDNA and RNA phages poses significant challenges to their detection using standard plaque assays or phage infection assays in liquid culture. As the preponderance of skin virome studies have focused on DNA phages, we encourage future investigations to employ multi-omic and expanded cultivation approaches to capture the full spectrum of *C. acnes* phage diversity, including RNA phages and those with cryptic or chronic life cycles.

Many studies have proposed phage therapy as a solution to cull specific *C. acnes* subpopulations on human skin^18,20,29–31^. In this work, we provide evidence that there are likely barriers to phage-mediated lysis on human skin that need to be considered when designing targeted microbiome-based therapeutics. Therefore, therapies leveraging naturally occurring *C. acnes* phage may have limited efficacy, and a limited ability to remodel the microbiome with strain-level precision, due the inefficiency of phage-mediated lysis on human skin.

The static arms race dynamics between *C. acnes* and its phage presented here represent an important counterexample to canonical models of antagonistic coevolution. We encourage future research to identify more examples of evolutionary stasis in virus-microbe dynamics across diverse taxa and ecosystems, to enable the study of the specific ecological conditions that favor the emergence of static arms races rather than escalating antagonistic coevolution.

## Materials and methods

### Material availability

This study did not generate new unique reagents

### Data and code availability

All original code has been deposited at GitHub and is publicly available as of the date of publication: https://github.com/adtripp/Tripp_et_al_cacnes_phage_defense. Any additional information required to reanalyze the data reported in this paper is available from the lead contact upon request.

### Figures and statistical analysis

Data processing and analysis were performed using pandas v1.3.4^36^ and R v4.1.3^37^, the latter of which was also used for all figure generation and statistical testing. Unless otherwise specified, comparisons between experimental and control groups were conducted using a two-sided Wilcoxon rank-sum test to assess differences in group medians.

### Bacterial genomes and phylogenetic analysis

To reconstruct the global diversity of *C. acnes*, we first curated 3032 whole-genome sequences from three sources: 2167 and 865 natural isolates previously cultured from human skin by our laboratory^11,12^, and 173 genomes labeled as ‘*Cutibacterium acnes*’ in the 661K bacterial genome database^38^. Raw sequencing reads were quality-filtered using Bracken v2.5^39^ to retain only those samples in which ≥ 90% of reads were taxonomically assigned to *C. acnes*. Filtered reads were assembled into draft genomes using SPAdes v3.13^40^ and functionally annotated with Prokka v4.8.1^41^. Assemblies with high fragmentation (i.e., composed of more than 200 contigs) were excluded from downstream analyses to ensure genomic integrity.

For phylogenetic reconstruction, genomes were dereplicated into distinct lineages using the reference-based single nucleotide variant (SNV) calling approach as described in prior studies^11,12,42^. Briefly, each genome was aligned to the Pacnes_C1 reference genome (*C. acnes* NC_018787), SNVs were identified in the core genome, and isolates were clustered based on pairwise core-genome mutational distances. Within each lineage, a single representative genome was selected based on the highest median sequencing coverage; singleton isolates (those not belonging to defined lineages) were retained. This resulted in a final set of 360 representative isolates used to construct the phylogenetic tree. A list of all genomes and corresponding phylogenetic information is available in Table S1.

### Defense system detection and defense island localization

To annotate the *C. acnes* pan-immune arsenal, anti-phage defense systems were identified across the curated genome dataset using DefenseFinder v1.0.8^32^ with default parameters. To evaluate whether defense systems were enriched in specific genomic regions -- potentially indicative of defense islands -- we analyzed their genomic context as follows: for each detected system, we extracted the contig containing the defense genes along with 5,000 bp of flanking sequence on both the upstream and downstream sides. These extended contigs were aligned to the Pacnes_C1 reference genome (*C. acnes* NC_018787) using BLAST v2.7.1^43^ under default settings. A defense system was classified as localized to a specific genomic region if ≥ 10% of the corresponding contigs mapped to the same locus in the reference genome. All defense systems detected by DefenseFinder^32^ are available in Table S2.

### Identification of incomplete and truncated defense systems

To assess the completeness of detected defense systems, we investigated the presence of defense genes across the *C. acnes* phylogeny and looked for cases of pseudogenization. We generated representative sequences for each defense gene family using CD-HIT v4.8^44^ (cd-hit-est; parameters: -c 0.95 -s 0.9 -T 0 -M 0 -d 0). These representative sequences were used to build a custom BLAST nucleotide database. Using BLAST v2.7.1^43^ with default parameters, we screened each genome assembly for homologs of the required defense genes. A gene was considered present if ≥ 95% of the representative sequence aligned to a region in the query genome. To minimize false positives due to assembly fragmentation, hits located within 500 bp of a contig end were excluded.

A defense system was classified as incomplete if one or more essential genes were missing from the genome. In addition, we examined gene annotations for evidence of truncation -- such as premature stop codons or frameshift mutations -- by comparing the length of predicted open reading frames (ORFs) to the representative full-length sequence. A gene was considered truncated if its coding sequence was at least 100 bp shorter than the reference. A defense system was deemed complete only if all mandatory components were both present and full-length.

### Defense system detection in skin-associated bacterial genera

To contextualize *C. acnes’* limited defense repertoire, we examined the distribution of anti-phage defense systems in other skin-associated bacteria. Defense system annotations were obtained from the DefenseFinder wiki Resource database v1.0^45^, a comprehensive resource for bacterial and archaeal defense systems derived from RefSeq genomes. Data were extracted for two major bacterial phyla: *Actinomycetota* and *Bacillota*. From these datasets, we focused our analysis on four skin-associated bacterial genera: *Corynebacterium, Cutibacterium, Staphylococcus*, and *Streptococcus*, which represent key components of the human skin microbiome and are relevant to skin health and disease. Summary statistics were calculated for each genus and species, including the number of isolates analyzed, minimum and maximum defense system counts, and median values. Statistical comparisons between genera were performed using the Wilcoxon rank-sum test. We performed additional pairwise comparisons between all *Cutibacterium* species present in the dataset using the same statistical approach.

### Gene gain and loss analysis

To test whether genome-wide evolutionary constraints explain the conserved defensome, we quantified gene turnover across the *C. acnes* phylogeny. We focused on the 93 *C. acnes* lineages defined by Baker et al.^12^, using the associated phylogenetic trees (as described in the original study). Only lineages containing more than ten isolates were included to ensure robust statistical inference. Gene families were clustered using Roary v3.13.0^46^ with default parameters. To reduce noise from potential contamination or assembly artifacts, gene clusters present in fewer than two isolates within a lineage or in less than 3% of isolates across the lineage were classified as absent.

For each gene cluster, ancestral states were reconstructed using PastML v1.9.15^47^ under the Joint prediction method and the F81 character evolution model. The number of gene gain and loss events was inferred by counting transitions between presence and absence states across internal branches of the phylogeny.

### Carriage of plasmids and prophages in *C. acnes* isolate genomes

To assess the mobility of plasmids and prophages, we screened all genome assemblies for known components of the *C. acnes* mobilome and annotated their presence in the context of the core genome phylogeny. The *C. acnes* mobilome consists of a linear tight adherence (TadE) plasmid^61–64^, a novelly identified circular rep-encoding single-stranded (pCRESS) plasmid^65^, and a prophage that exists as a non-integrative pseudolysogen. Raw sequencing reads were mapped to reference sequences for each known mobile genetic element using bowtie2 v2.2.6^50^ with parameters: -X 2000 --no-mixed --dovetail. An element was considered present if it was detected at ≥ 1X vertical coverage (mean depth) and ≥ 50% horizontal coverage (breadth). To identify potential integration sites, we aligned plasmid and prophage sequences to the Pacnes_C1 reference genome (*C. acnes* NC_018787) using BLAST v2.7.1^43^. Integration loci were determined through manual inspection of alignment coordinates. An element was considered integrated into a specific genomic region if ≥ 10% of contigs carrying the element overlapped the same locus in the reference genome.

### CRISPR-Cas Array and Spacer Analysis

To explore adaptive immunity in *C. acnes*, we identified CRISPR-Cas arrays and analyzed spacer content. CRISPR-Cas arrays were identified using CCTK v1.0.2^34^. We ran CCTK on all genome assemblies to detect CRISPR arrays and to identify array repeats specific to *C. acnes*. Next, to improve sensitivity and capture spacer diversity, particularly in repetitive regions poorly resolved in draft assemblies, we applied SpacerExtractor v1.0^35^ directly to raw short-read sequencing data from *C. acnes* phylogroup K isolates. Spacer sequences were then dereplicated using CD-HIT v4.8^44^ (cd-hit-est; parameters: -c 1 -s 1 -T 0 -M 0 -d 0) to remove exact duplicates. All CRISPR Cas array spacers detected by SpacerExtractor^35^ are available in Table S4.

To identify potential targets of spacers, all unique spacers were aligned to two databases using BLAST v2.7.1^43^: (1) a custom database of *C. acnes*-associated mobile genetic elements (including plasmids and prophages detected in this study), and (2) the IMG/VR and IMG/PR databases, which contain viral and plasmid sequences. Up to two nucleotide mismatches were permitted in alignments to allow for natural sequence variation.

### Liquid growth curve phage infection assay

To functionally test phage resistance, we developed a high-throughput liquid growth assay. From glycerol freezer stocks, isolates were streaked onto Reinforced Clostridium Medium (RCM) agar plates and incubated for 5 days to obtain single colonies. A single colony was inoculated into 500 µL of liquid RCM and grown for 48 hours to reach mid-log phase. Cultures were then diluted to an initial optical density at 600 nm (OD_600_) of 0.01 in 150 µL of fresh medium and transferred to 96-well microtiter plates (Greiner BioOne). Phage or control (RCM) was added at a multiplicity of infection (MOI) of 0.001. Cultures were incubated at 33°C with shaking at 250 rpm for 72 hours in a LogPhase 600 plate reader (Agilent), with OD_600_ measurements recorded every 20 minutes. Plates were sealed with Breath-Easy sealing film (Research Products International, Cat. No. 248738) to prevent evaporation. Each condition was performed in quadruplicate, and experiments included both phage-free and cell-free negative controls.

A total of 109 bacterial isolates, representing all major *C. acnes* clades (Figure S6), were challenged with a *C. acnes* phage cocktail. To quantify phage-induced growth inhibition, we calculated an inhibition score based on the area under the growth curve (AUC), adapted from a previously described method^48^. First, background-subtracted OD_600_ values were obtained by subtracting the initial OD from all time points. The AUC was computed for both phage-treated and untreated (control) cultures grown on the same microtiter plate. The inhibition score was defined as the mean AUC of phage-treated replicates, expressed as a percentage of the mean AUC of the corresponding phage-free control. Lower scores indicate stronger phage-mediated inhibition. Data are in Table S3.

### Healthy facial skin metagenome dataset

To assess ecological patterns of phage and host in natural populations, we compiled a global metagenomic dataset. We searched the NCBI SRA database for publicly available human facial skin metagenomes using the keywords ‘skin metagenome’ and ‘human skin metagenome’ (accessed December 2023). Studies were excluded if they had fewer than 10 subjects, did not include subject identifiers, or sampled subjects that had recently taken antibiotics or other skin therapeutics. In addition, studies were removed if they did not sample from facial skin, and if they had fewer than a million reads. We excluded subjects from the dataset with acne or other skin diseases according to the metadata provided by each study.

Metagenomic sequencing data was dereplicated by subject as follows: for studies that collected samples at multiple timepoints per subject, the time point with the highest sequencing depth was chosen. For studies that collected samples at multiple facial sites per subject, the facial site with the highest average sequencing depth for each study was chosen, followed by the time point with the highest sequencing depth if applicable. For studies that collected multiple samples per subject using different sampling methods, we chose samples that were collected via facial swab to standardize method consistency with the rest of the dataset. After this quality control of the initial 621 samples, the final dataset comprised samples from 471 individuals from 3 continents across 7 independent studies. A list of all metagenomes and corresponding metadata is available in Table S5.

### PHLAME metagenomic analysis of strain-level diversity

To estimate *C. acnes* subphylogroup relative abundances in metagenomic samples, we applied PHLAME v1.0^13^, a phylogeny-aware probabilistic tool for strain deconvolution. Raw metagenomic reads were first quality-trimmed using sickle v1.33^49^, then aligned to the Pacnes_C1 reference genome (*C. acnes* NC_018787) using bowtie2 v2.2.6^50^ with parameters: -X 2000 --no-mixed --dovetail. Duplicate reads were removed using SAMtools markdup v1.15.1^51^. Only samples achieving a median coverage of at least 5X across the *C. acnes* genome were retained for downstream analysis, ensuring sufficient bacterial abundance.

Aligned reads were classified using PHLAME with default parameters (-n 0.1 -p 0.1) and a custom reference database containing the 360 representative *C. acnes* genomes described above. Clades -- representing intraspecies diversity -- were defined at the subphylogroup level using established typing schemes for *C. acnes*^13,52^ (as implemented in Qu et al.). A subphylogroup was considered detected if its estimated relative abundance exceeded 1% of the total *C. acnes* population in the sample.

### Metagenomic analysis of prevalence and virus-to-microbe ratios

To quantify phage abundance relative to host, we calculated metagenome-derived virus-to-microbe ratios (mVMR). We assessed the presence of *C. acnes* phage by mapping metagenomic reads to the reference phage genome NC_028967.1 (a representative *C. acnes* siphovirus) using bowtie2 v2.2.6^50^ with the same alignment parameters as above. A sample was considered phage-positive if at least 10% of the phage genome was horizontally covered at 1X depth. The mVMR was calculated as the ratio of *C. acnes* phage reads normalized by phage genome length (in kb) to bacterial *C. acnes* reads normalized by bacterial genome length (in kb). This metric provides an estimate of phage abundance relative to host bacteria within each sample.

## Acknowledgments

We thank the participants of the original study, the MIT BioMicroCenter for assistance in sequencing, and all members of the Lieberman Lab for advice on this project and feedback on the manuscript. This work was funded by NIH grants 1DP2GM140922 and 1R01AR084097 (to TDL), the National Science Foundation Graduate Research Fellowship under Grant No. 2139433 (to ADT), and the U.S. Department of Energy Office of Science, Workforce Development for Teachers and Scientists, and Science Graduate Student Research (SCGSR) program under award number DE-SC0014664 (to ADT). The work conducted by the U.S. Department of Energy Joint Genome Institute (https://ror.org/04xm1d337), a DOE Office of Science User Facility, which is supported by the Office of Science of the U.S. Department of Energy operated under Contract No. DE-AC02-05CH11231. This work was performed in part at Aspen Center for Physics, which is supported by National Science Foundation grant PHY-2210452

## Author contributions

Conceptualization: ADT, TDL; Methodology: ADT, TDL; Investigation: ADT, EQ, IB, JB, JSB, CPM, TDL; Funding acquisition: TDL; Writing – original draft: ADT; Writing – review and editing: ADT, JB, CPM, SR, FAM, TDL.

