## Supplemental Figures 1-13 for "Phage-mediated lysis does not determine *Cutibacterium acnes* colonization on human skin"

**BioRxiv, 2025-09**

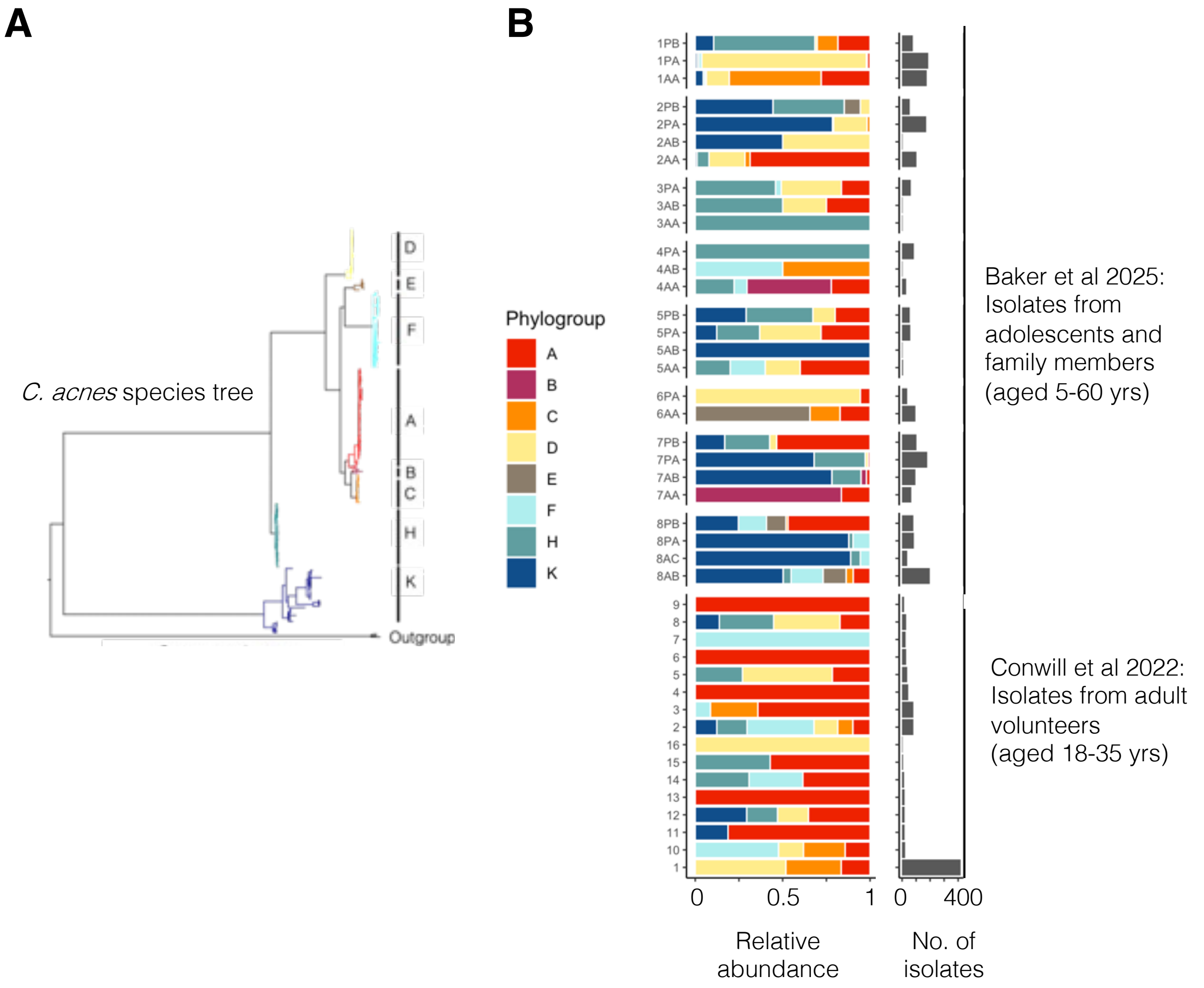


**Figure S1. Multiple distinct strain types of *C. acnes* co-exist on individuals' skin.** (A) Core-genome reference phylogeny of *C. acnes* constructed from 360 representative genomes. Major intraspecies clades (phylogroups) are indicated by letter and color coded (Methods). (B) Reanalysis of published culture-based whole-genome sequencing data from Baker et al., 2025 [12] and Conwill et al., 2022 [11]. This data comprises 3,032 *C. acnes* genomes previously sampled by our lab from the sebaceous skin of 43 different individuals aged 5 to 60 years old (Table S1). Bar plot showing the relative abundance of different phylogroups amongst the set of sequenced isolates from subjects with at least 10 isolates. On the majority of subjects, it is common for multiple phylogroups of *C. acnes* to co-exist.

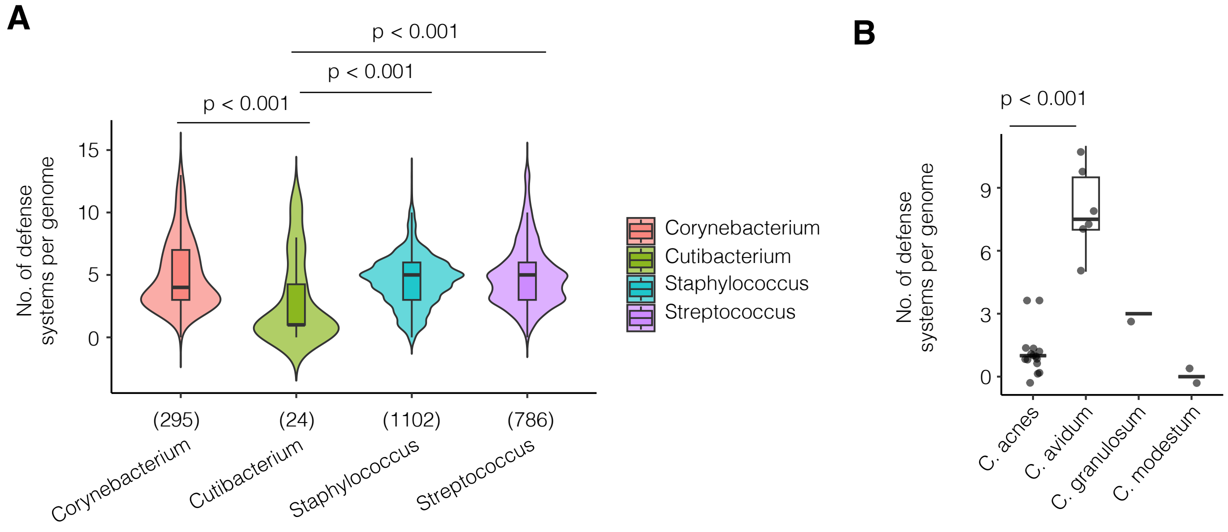


**Figure S2. *Cutibacterium* have fewer anti-phage defense systems per genome compared to other genera commonly found on human skin, related to Figure 1.** (A) Violin plot of the number of defense systems per genome for four common genera of bacteria found on human skin, as reported in the DefenseFinder Wiki database v1.0 [45] (Methods). Despite similar genomes sizes (~2.5 Mb), on average bacterial genomes from the genus *Cutibacterium* have significantly fewer defenses compared to genomes from the genus *Corynebacterium, Staphylococcus* and *Streptococcus* (for all comparisons, Wilcoxon rank-sum test, $p < 0.001$). Mean number of defense systems in each genus: *Cutibacterium* 2.9; *Corynebacterium* 5*;* *Staphylococcus* 4.5; *Streptococcus* 4.9. (B) Boxplot and jitter of the number of defense systems per genome within the genus *Cutibacterium*. Notably, *C. avidum* stands as a statistical outlier with significantly more defenses per genome compared to *C. acnes* (Wilcoxon rank-sum test, $p < 0.001$).

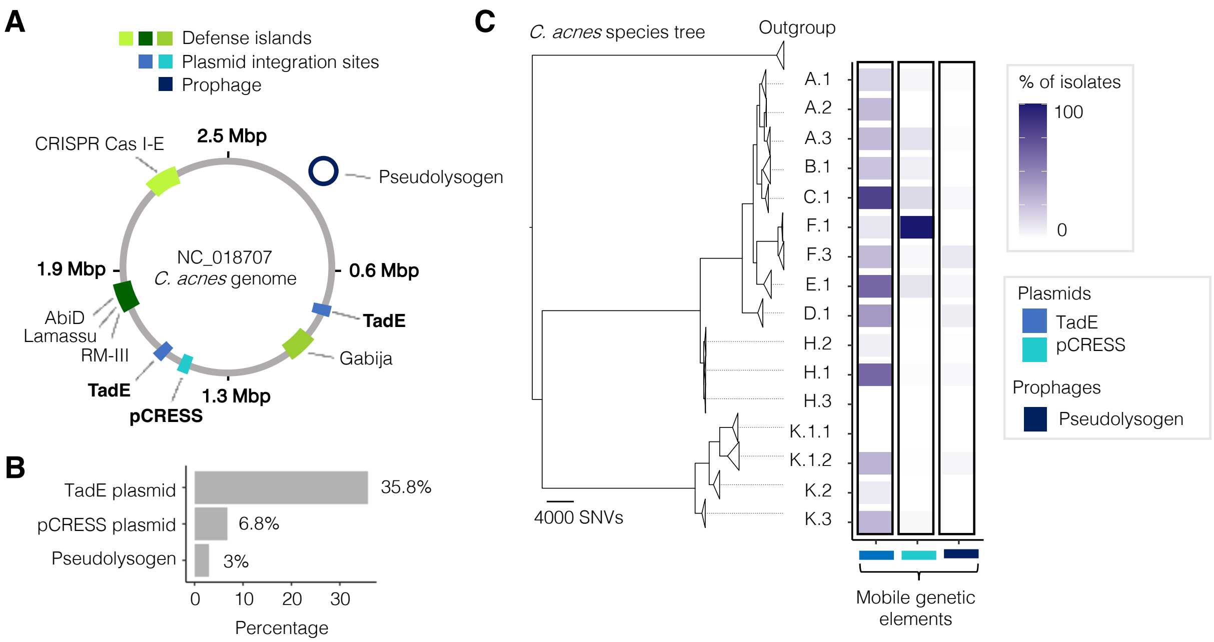


**Figure S3. Comparison of the mobility of defense islands in *C. acnes* to other mobile genetic elements, related to Figure 1.** (A) Genome map of the known *C. acnes* mobilome, which consists of three defense islands, two integrative plasmids (TadE and a novelly reported pCRESS), and a prophage that exist as a non-integrative pseudolysogen. (B) Bar chart of the percentage of isolates that carry the TadE and pCRESS plasmids and the pseudolysogen (Methods). (C) A maximum-likelihood phylogenetic tree constructed at the subphylogroup level shows the carriage of these mobile genetic elements (Methods). The two integrative plasmids and pseudolysogen have high levels of mobility, as evident by their sparse carriage throughout the phylogeny.

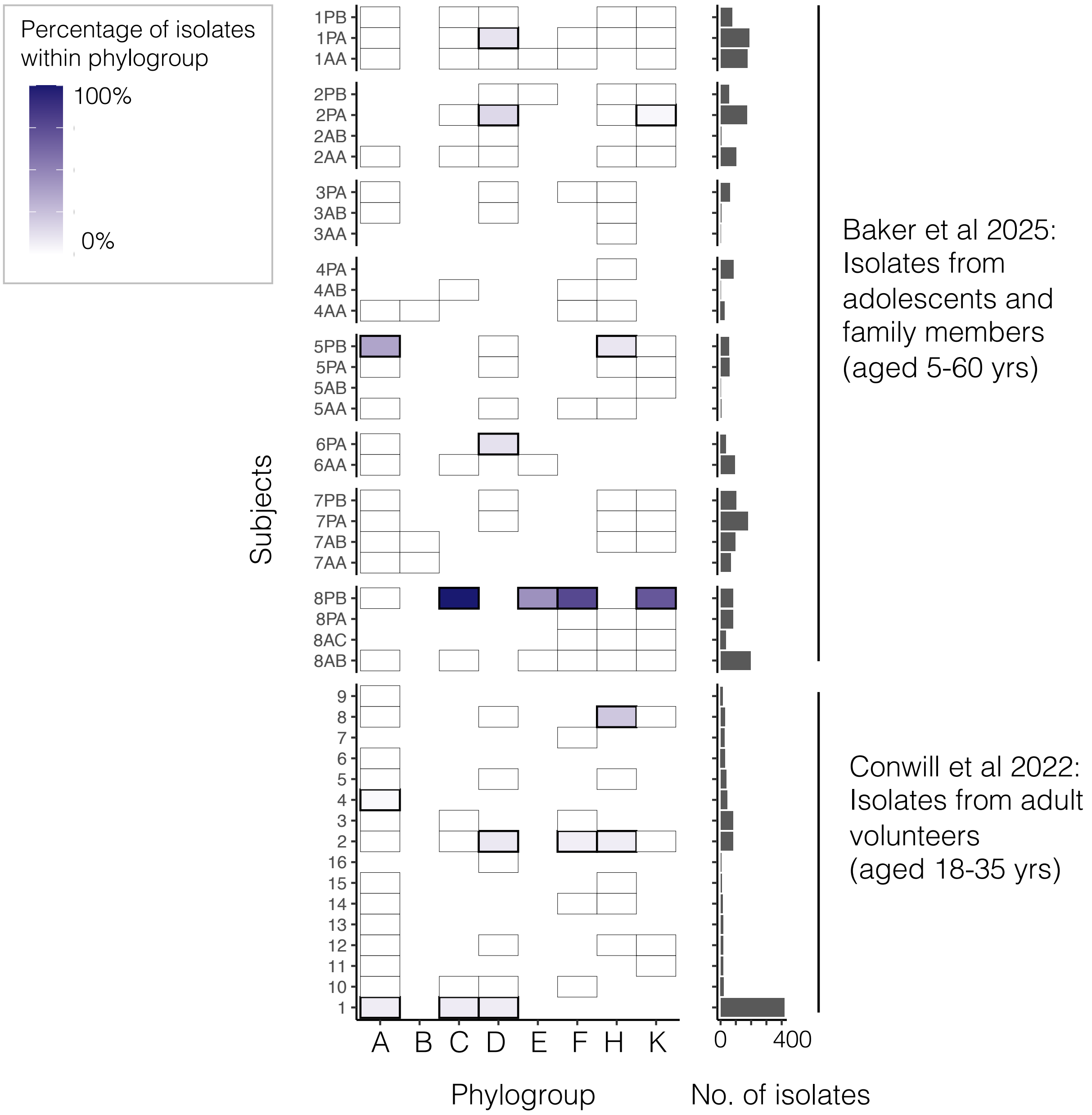


**Figures S4. Pseudolysogen carriage is rare and found sparsely across individuals.** Analysis of published culture-based whole-genome sequencing data from Baker et al., 2025 [12] and Conwill et al., 2022 [11] (Table S1). To examine pseudolysogen carriage across phylogroups and subjects, we plotted a heatmap showing the percentage of pseudolysogen-containing isolates within a phylogroup by subject. We found that more than half of pseudolysogen-containing isolates (57.1%) originated from just two subjects, suggesting potential sample bias or high inter-individual variation. Collectively, only 3% of 3,205 sequenced *C. acnes* isolates carried pseudolysogens.

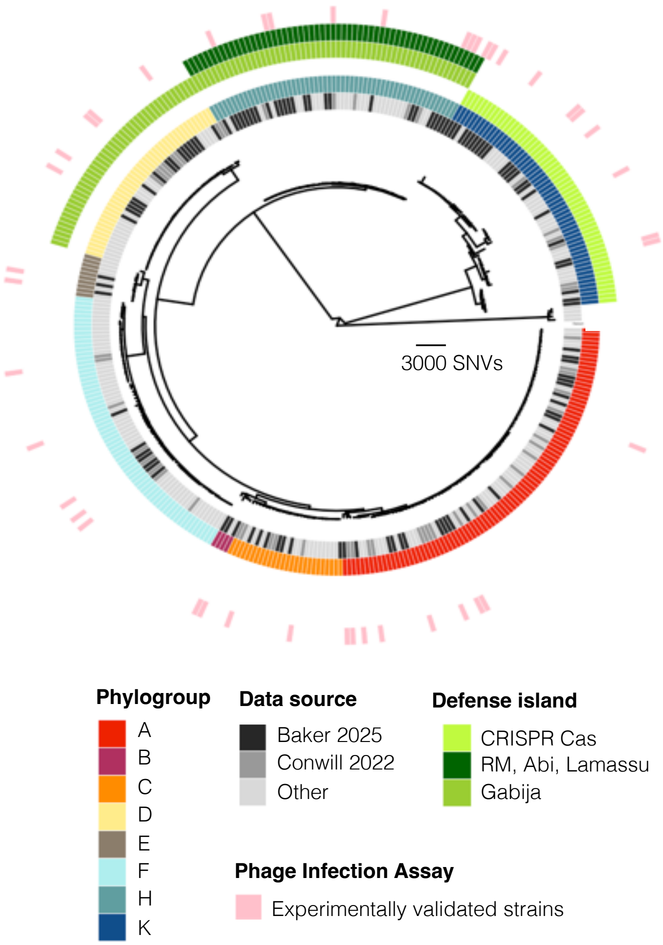


**Figure S5. Core-genome reference phylogeny of *C. acnes* constructed from 360 representative genomes, related to Figure 2.** Tree annotated with metadata of isolate phylogroup, data source, and defense island carriage (Table S1 and S2). Tips annotated in pink denote lineages from which sets of isolates were experimentally validated for susceptibility to phage infection (Figure 2). A total of 109 isolates representing *C. acnes* diversity at multiple evolutionary scales were experimentally tested for susceptibility to infection by a phage cocktail (Methods).

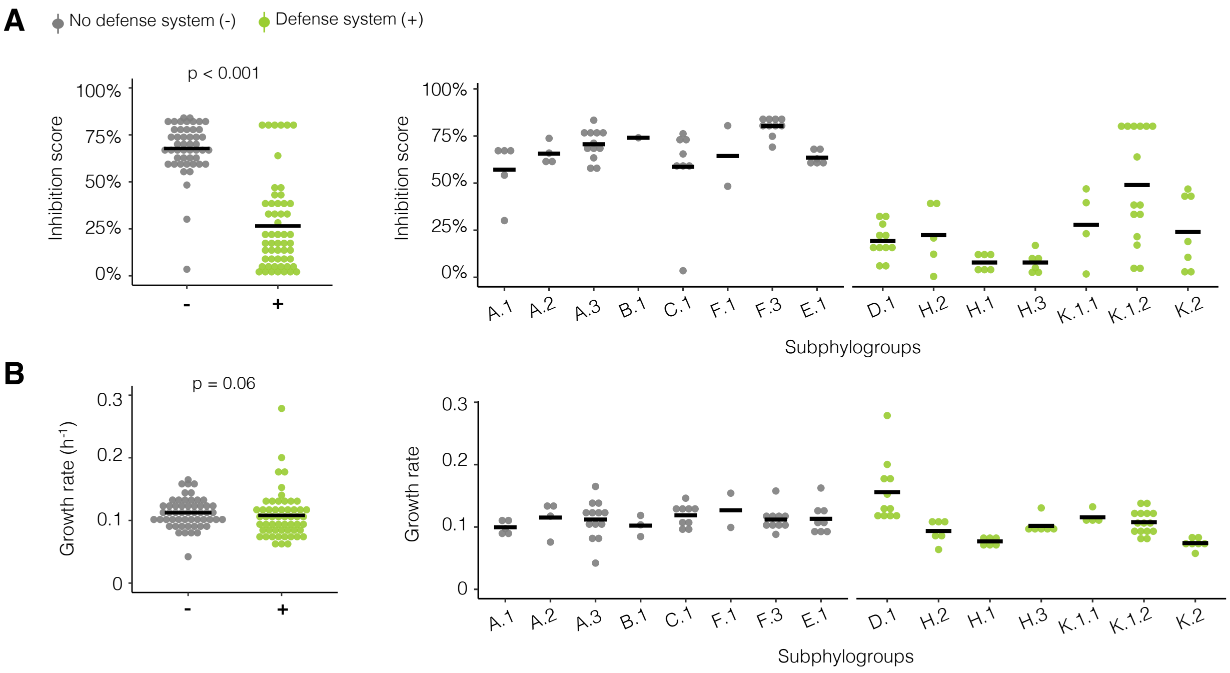


**Figure S6. Experimental assessment of *C. acnes* inhibition score and growth rate, related to Figure 2.** (A) Plot of the inhibition scores calculated for the 109 *C. acnes* isolates infected in liquid culture with our phage cocktail (Figure S5). Inhibition scores are calculated as the ratio of the Area Under the Curve (AUC) for untreated bacterial cultures versus phage-treated cultures (Table S3; Methods). The left dot plot aggregates data and plots the mean value for each subphylogroup. Right dot plot aggregates data by isolate and plots the mean value for each isolate, organized by subphylogroup. Each isolate was grown in biological triplicates. Isolates lacking defense systems have significantly higher inhibition scores compared to those that carry defense systems, indicating greater susceptibility to phage predation (Wilcoxon rank-sum test, $p < 0.001$). Notably, the highest observed inhibition score was 84.2%, as some bacterial growth always occurred before phage lysis initiated. (B) To detect fitness costs associated with defense system carriage we also measured the growth rates ($\mu = h^{-1}$). Isolates that carry or lack defense systems revealed no significant difference in growth rates (Wilcoxon rank-sum test, $p = 0.19$).

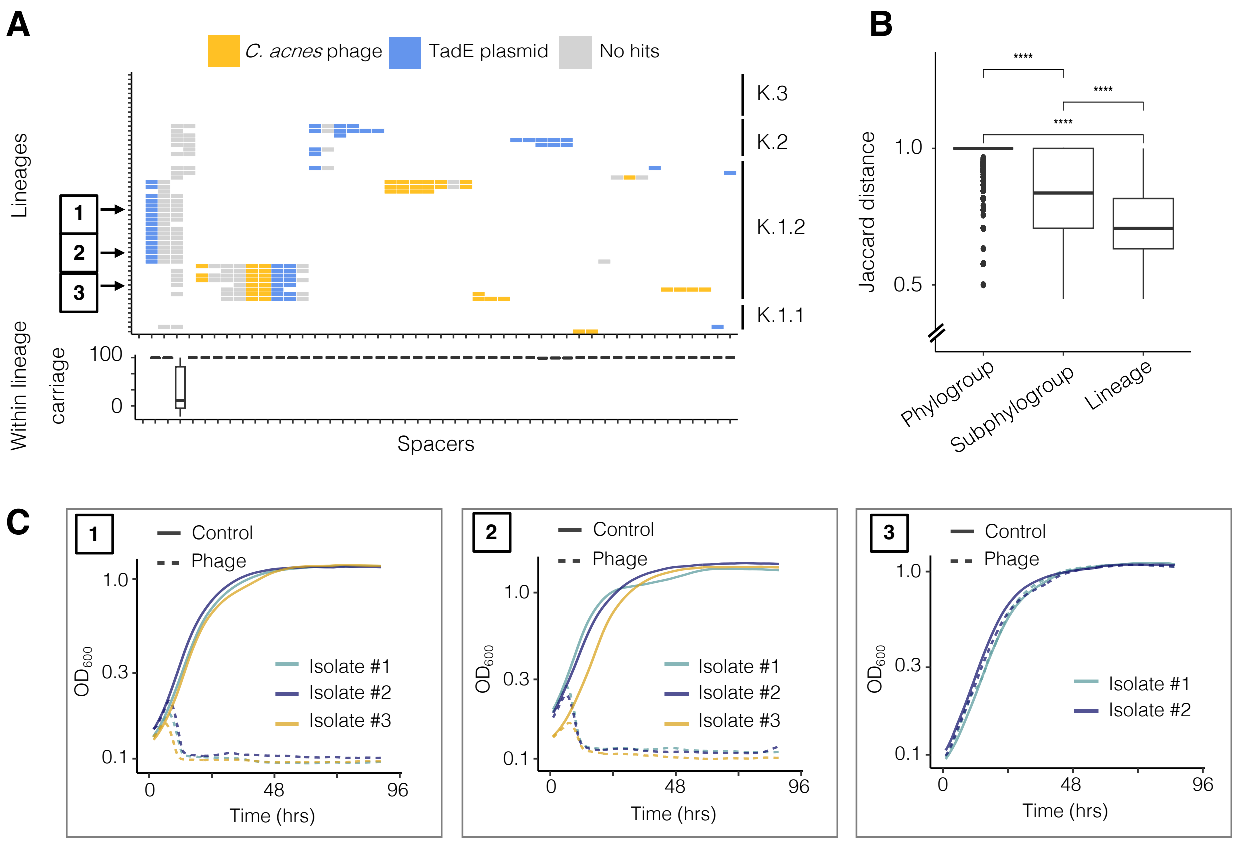


**Figure S7. Conservation of *C. acnes* CRISPR Cas array spacer content at the lineage-level and turnover on longer evolutionary scales, related to Figure 2.** (A) To investigate if the variation within isolates from phylogroup K can be explained from the content of spacers in their CRISPR Cas Type I-E arrays, we examined spacer targets in the context of the core phylogeny (Table S4; Methods). *C. acnes* phylogroup K genomes were screened for CRISPR Cas arrays, and the spacers were extracted and clustered to within 3 SNPs. Spacers were blasted against a database of *C. acnes* mobile genetic elements to identify targets. We identified spacers that target *C. acnes* phage in the CRISPR Cas arrays of some, but not all genomes. (B) We noted divergence in *C. acnes* CRISPR Cas array spacer content across evolutionary scales (Wilcoxon rank-sum test, $p < 0.001$ for all comparisons). CRISPR Cas array spacer content between pairs of isolates from the same lineage have significantly lower Jaccard distances compared to those from higher order taxonomic groupings (i.e., subphylogroups and phylogroups). This analysis shows a lack of complete conservation and some turnover as time goes on (beyond lineage level). (C) Growth curves of representative isolates from three different lineages in subphylogroup K.1.2 illustrate phenotypic conservation of bacterial susceptibility to phage within lineages (Methods).

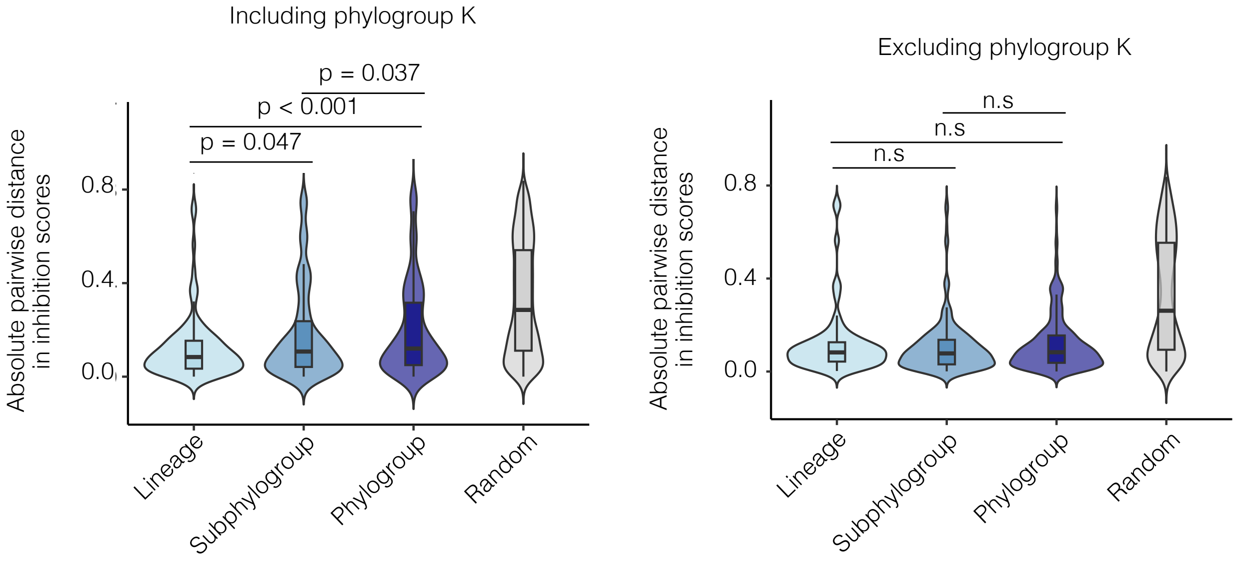


**Figure S8. Inclusion of phylogroup K adds significant heterogeneity in inhibition scores across evolutionary timescales, related to Figure 2.** We compared pairwise distances in inhibition scores across different phylogenetic groupings (lineage, subphylogroup, and phylogroup) (Figure S5 and S6; Table S3). Violin plot of absolute pairwise distance in inhibition scores compared within phylogenetic groups based on the inclusion or exclusion of isolates from phylogroup K. When we analyzed the data excluding isolates from phylogroup K, we found no significant differences in pairwise distances across any phylogenetic level (Wilcoxon rank-sum test, all comparisons $p > 0.1$), suggesting that the degree of variation in inhibition scores is the same across evolutionary scales. As expected, reintroducing phylogroup K isolates added significant heterogeneity due to this phylogroup encoding a CRISPR Cas type I-E system known to confer spacer-dependent anti-phage activity (Figure S7; Table S4).

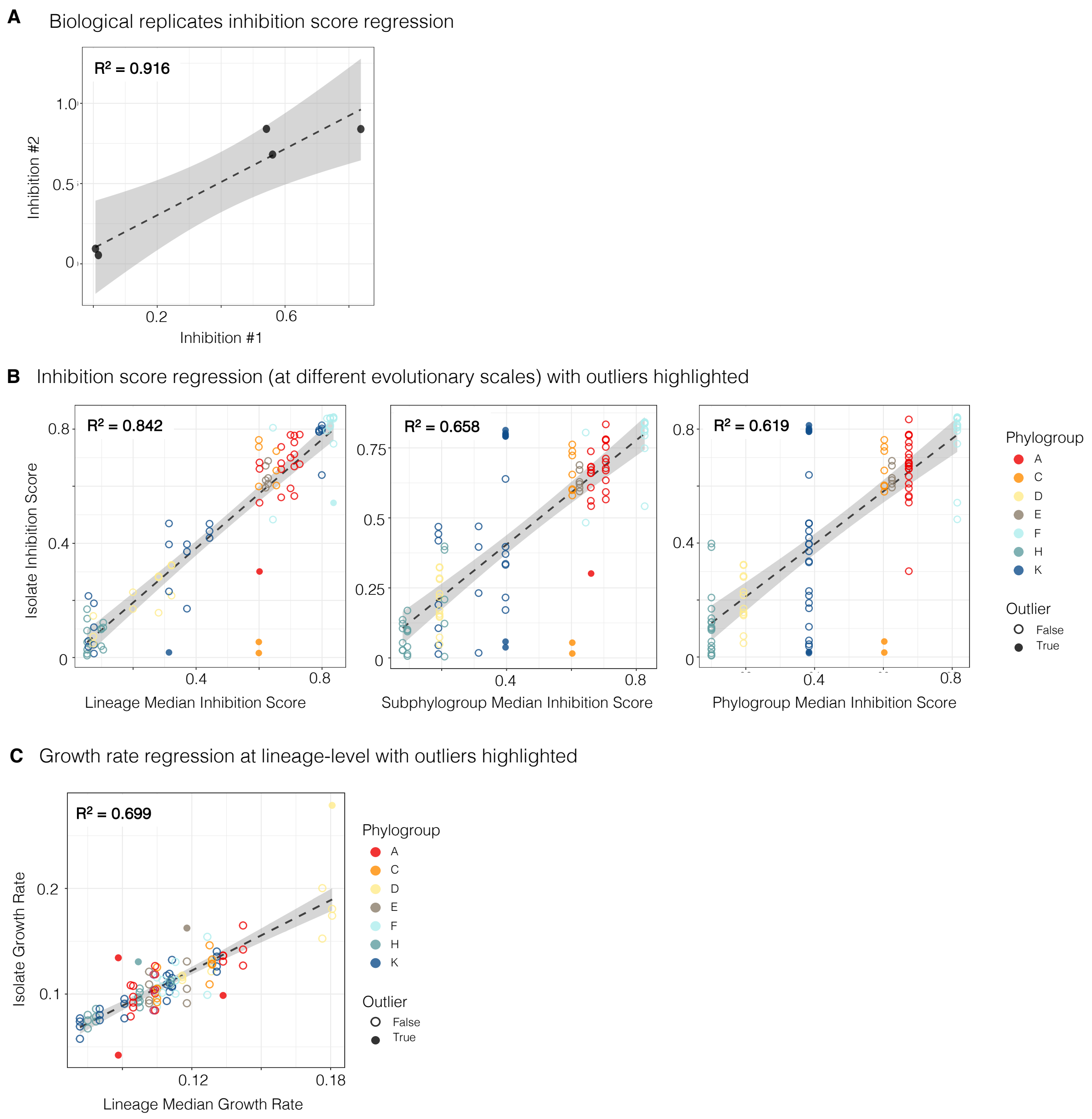


**Figure S9. Evolutionary conservation of phage inhibition scores and bacterial growth rate, related to Figure 2.** (A) Correlations between inhibition scores for biological replicates. (B) Correlations between isolate inhibition scores and the median inhibition scores of their respective phylogenetic group across taxonomic levels, with regression lines (grey) and outliers highlighted (closed circles). Scatterplots show squared correlation coefficients ($R^2$) for phylogroups, subphylogroups, and lineages (each dot represents n = 109 isolates colored by phylogroup; Figure S5), corresponding to Figure 2B and 2C. (C) Lineage-level growth rate variability. Scatterplot of individual isolate growth rates versus their respective lineage medians, with outliers (n = 6) identified in phylogroups A (n = 3), D, E, and H. Outliers were defined as isolates with standardized residuals > 2 in absolute value from the respective linear regression model. Data available in Table S3.

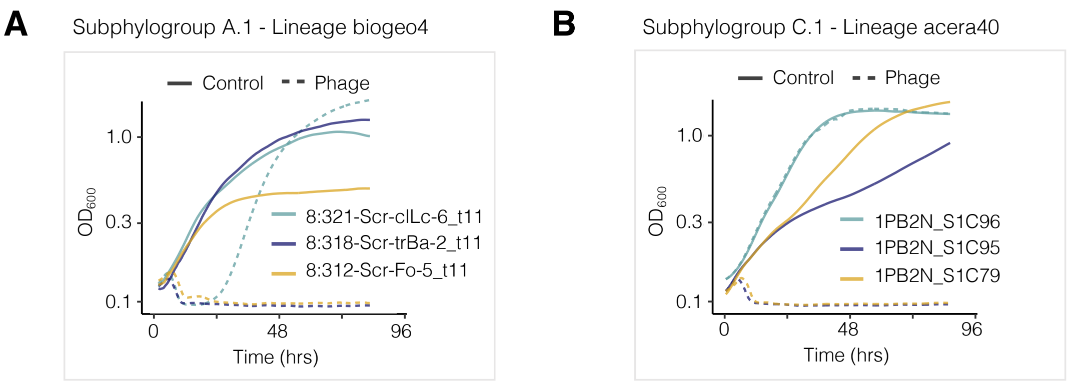


**Figure S10. Two cases of rapid resistance evolution in isolates lacking defense systems, related to Figure 2.** (A-B) Phage infection assay growth curve plots (Table S3; Methods), corresponding to Figure 2C outlier points. Solid line represents the un-treated culture (control) and dashed line represents the phage-treated culture. Isolates were grown in triplicates, and a single replicate is shown here. There were two cases of isolates that lacked defense systems but exhibited low inhibition scores (resistance to phage infection). One of these isolates evolved resistance during the experiment (Isolate A.1 8:321-Scr-clLc-6_t11; biogeo4; green line), and the other appears to be the sole case of natural occurring resistance in an isolate lacking any defense system (Isolate C.1 1PB2N_S1C96; acera40; green line). In both cases, two other isolates from these lineages, sampled from the same subject at the same timepoint, remained susceptible to phage infection.

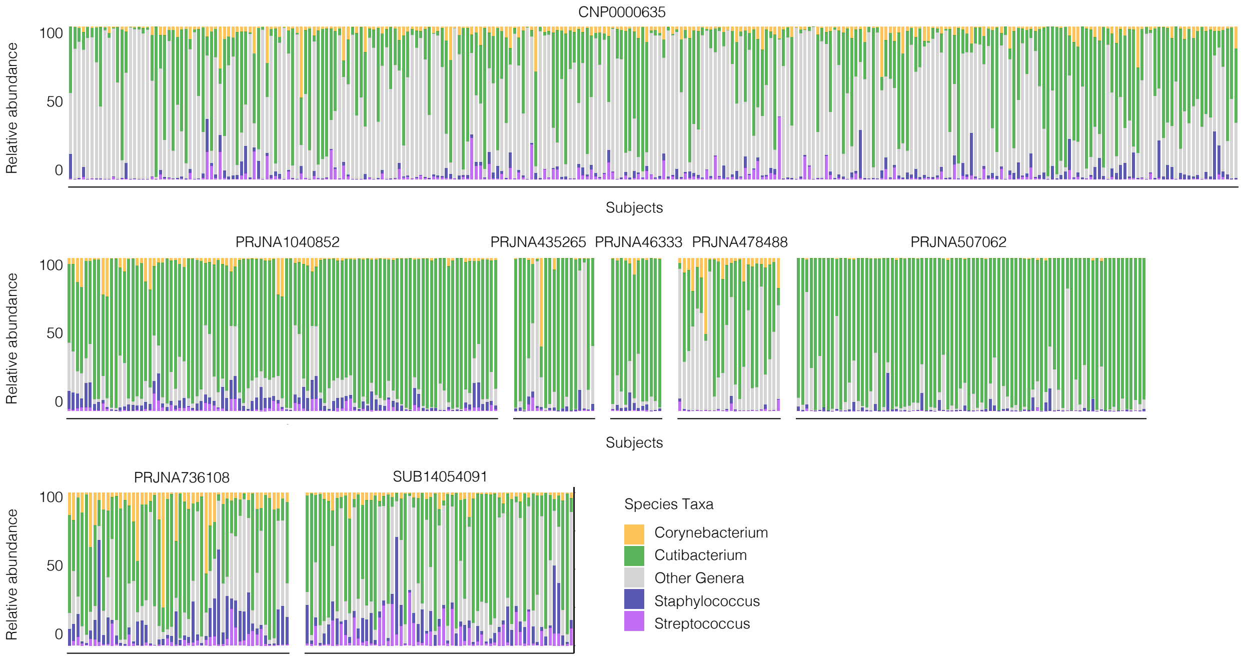


**Figure S11. On-person taxonomic abundances of major skin taxa, related to Figure 3 and 4.** Bar plots of the relative abundance of four major skin-associated genera (*Corynebacterium, Cutibacterium, Staphylococcus,* and *Streptococcus*) across the 621 publicly available human facial skin metagenomes from 3 continents across 8 independent studies analyzed in this work (Methods).

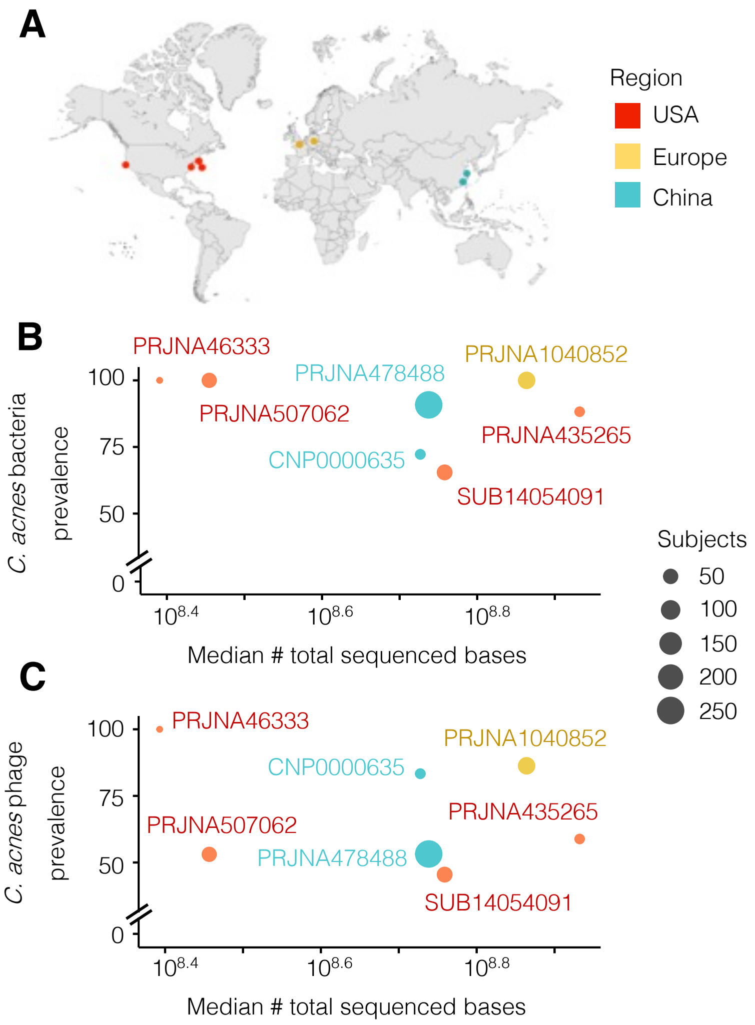


**Figure S12. Prevalence of *C. acnes* and its phage in global facial skin metagenomes, related to Figure 3 and 4.** (A) Geographic locations of sampling sites for metagenome meta-analysis (Created in https://BioRender.com). We analyzed 621 publicly available human facial skin metagenomes from 3 continents across 8 independent studies originating in the United States of America (red), Europe (yellow), and China (blue) (Table S5). Each region contained data from at least two independent research groups. After quality control filtering (Methods), the final dataset comprised samples from 471 individuals. (B-C) Relationship between bacteria and phage detection rates and sequencing depth. The scatter plot shows the percentage of individuals with detectable *C. acnes* bacteria (B) and phage (C) versus the median number of total sequenced reads (log10 scale) for each study. Data points are color-coded by region and labeled with the Bioproject. *C. acnes* bacteria was considered present if the NC_018787 reference genome was detected at 5X coverage. *C. acnes* phage was considered present if at least 25% horizontal coverage of the NC_028967.1 reference genome was detected at 1X coverage.

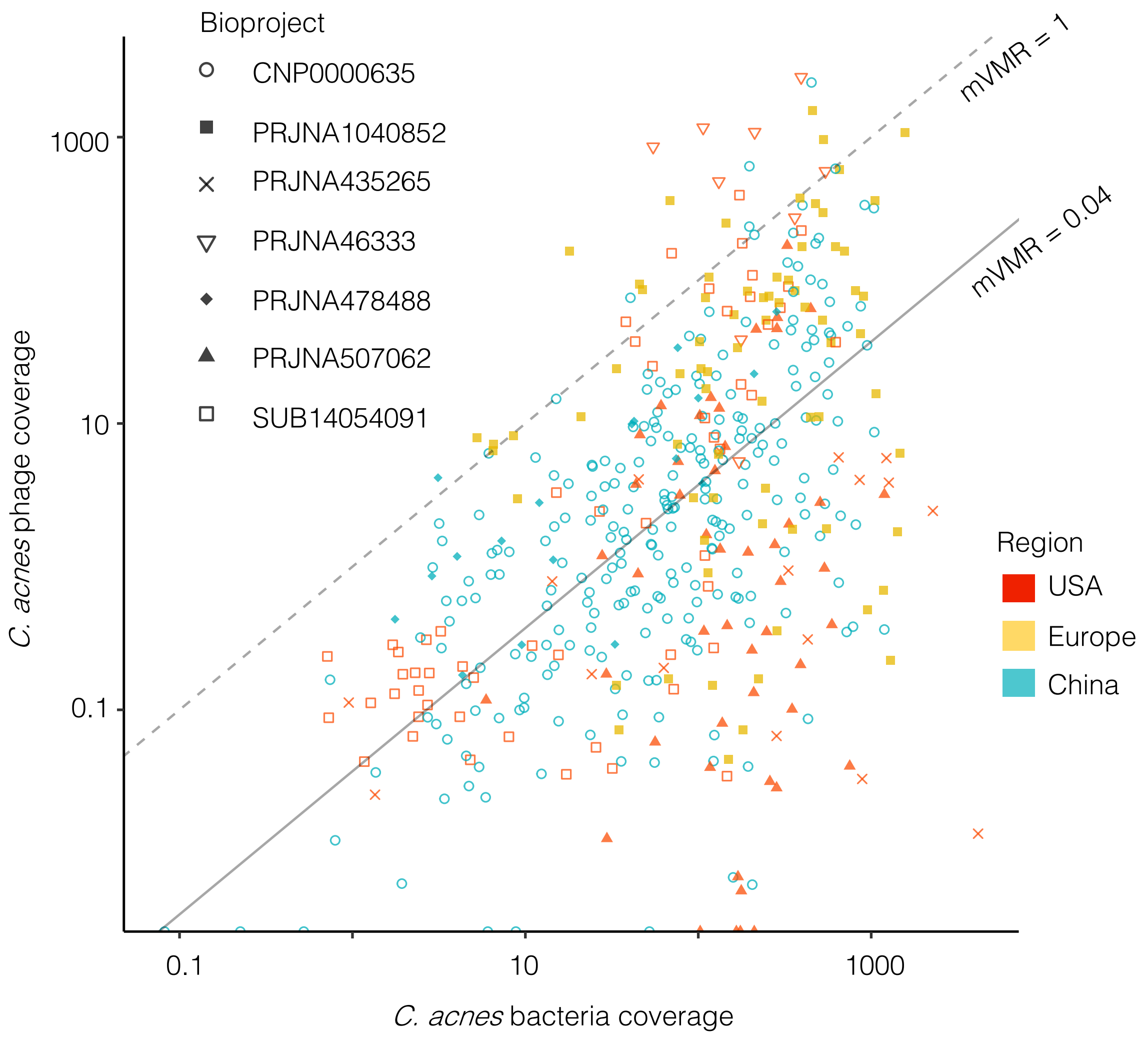


**Figure S13. Low *C. acnes* virus-to-microbe ratios in global facial skin metagenomes, related to Figure 4.** The scatter plot illustrates the relationship between the mean depth of coverage of the *C. acnes* bacterial versus phage genome (both on log10 scales) across different studies (Table S5; Methods). Each dot represents a single individual's metagenome, colored by study region, and the distinct shapes indicate from which Bioproject a sample originated. The dashed diagonal line represents a 1:1 and 0.04 metagenome-derived virus-to-microbe ratio (mVMR), serving as a reference for phage pressure intensity. In the majority of samples, the mVMR was low -- approximately 1 phage genome per 25 bacterial genomes (median mVMR = 0.04). Given the absence of lysogens and low prevalence of pseudolysogens (detected in < 3% of isolates, Figure S3), *C. acnes'* low mVMR suggests intrinsically limited lytic activity on human skin.
